# TGFβ2 regulates human trabecular meshwork cell contractility via ERK and ROCK pathways with distinct signaling crosstalk dependent on the culture substrate

**DOI:** 10.1101/2021.07.01.450718

**Authors:** Haiyan Li, Jessica L. Henty-Ridilla, Preethi S. Ganapathy, Samuel Herberg

## Abstract

Transforming growth factor beta 2 (TGFβ2) is a major contributor to the pathologic changes occurring in human trabecular meshwork (HTM) cells in primary open-angle glaucoma. Receptor binding of TGFβ2 activates non-canonical extracellular-signal-regulated kinase (ERK) and Rho-associated kinase (ROCK) signaling pathways, both broadly affecting HTM cell behavior. However, exactly how these signaling pathways converge to regulate pathologic HTM cell contractility associated with glaucomatous dysfunction is unclear. Here, we investigated the molecular mechanism underlying TGFβ2-induced pathologic HTM cell contractility, and the crosstalk between ERK and ROCK signaling pathways. We compared soft biomimetic hydrogels composed of collagen type I, elastin-like polypeptide, and hyaluronic acid with conventional stiff glass coverslips. Results show that HTM cell morphology and filamentous (F)-actin organization was affected by the underlying culture substrate: TGFβ2 increased HTM cell contractility via ERK and ROCK signaling pathways by differentially regulating F-actin, α-smooth muscle actin, fibronectin, and phospho-myosin light chain in cells grown on hydrogels compared to glass. Importantly, we showed that ERK inhibition further increased TGFβ2-induced phospho-myosin light chain levels in HTM cells on hydrogels, but not on glass, which translated into hypercontractility of three-dimensional (3D) HTM cell-laden hydrogels. ROCK inhibition had precisely the opposite effects and potently relaxed the TGFβ2-induced hydrogels. This suggests that ERK signaling negatively regulates ROCK-mediated HTM cell contractility, and that impairment of this crosstalk balance contributes to the pathologic contraction associated with the glaucomatous stressor TGFβ2. These findings emphasize the critical importance of using 3D tissue-mimetic extracellular matrix substrates for investigating HTM cell physiology and glaucomatous pathophysiology *in vitro*.

## 1. Introduction

Transforming growth factor beta 2 (TGFβ2), the predominant TGFβ isoform in the eye and aqueous humor, has been identified as a major player in contributing to the pathologic changes in primary open-angle glaucoma (POAG) with elevated intraocular pressure (IOP) (Agarwal et al., 2015; Fuchshofer and Tamm, 2009; Granstein et al., 1990; Inatani et al., 2001; Kasetti et al., 2018; Quigley, 1993). TGFβ2 is a key regulator of human trabecular meshwork (HTM) cell extracellular matrix (ECM) synthesis, deposition and degradation (Sethi et al., 2011). TGFβ2 has been shown to induce the synthesis/secretion of ECM proteins such as collagen type I and VI, laminin, elastin, fibronectin, plasminogen activator inhibitor 1 and myocilin, and increase the expression/activity of ECM crosslinking enzymes such as tissue transglutaminase 2, which irreversibly crosslinks fibronectin (Fleenor et al., 2006; Fuchshofer and Tamm, 2012; Fuchshofer et al., 2003; Han et al., 2011; Hann and Fautsch, 2011; Tamm et al., 1999; Tovar-Vidales et al., 2011; Welge-Lüßen et al., 2000). Furthermore, exposure of HTM cells to TGFβ2 induces the formation of F-actin stress fibers and expression of α-smooth muscle actin (αSMA), both strongly associated with cell contractility (Han et al., 2011; Pattabiraman and Rao, 2010).

Binding of active TGFβ2 to its receptor on the cell surface activates canonical Smad (Smad2/3) and diverse non-Smad signaling pathways including extracellular-signal-regulated kinase (ERK) and Rho-associated kinase (ROCK) (Ma et al., 2020; Prendes et al., 2013; Zhang, 2009). It has been shown that ERK inhibition in HTM cells decreased the expression of αSMA and fibronectin (Pattabiraman and Rao, 2010). Previous studies have also demonstrated that ROCK signaling in HTM cells is involved in regulating aqueous humor outflow resistance and thereby IOP homeostasis (Honjo et al., 2001; Rao et al., 2001). In the United States, only the recently approved ROCK inhibitor Netarsudil (Serle et al., 2018) directly targets the HTM to increase outflow via cell/tissue relaxation (Rao et al., 2001; Rao et al., 2017; Ren et al., 2016; Tanna and Johnson, 2018; Wang and Chang, 2014; Zhang et al., 2012). It has been shown that ROCK inhibition in HTM cells can potently reduce downstream phospho-myosin light chain (p-MLC) levels and assembly of actin stress fibers, jointly contributing to decreased HTM cell contractility (Pattabiraman and Rao, 2010; Ramachandran et al., 2011). Thus, both ERK and ROCK signaling are considered critical in regulating outflow and IOP; however, the required interplay between these signaling pathways underlying HTM cell behaviors remains unclear.

Most studies investigating HTM cell (patho)physiology to date have used conventional twodimensional (2D) cell monolayer cultures on tissue culture plastic or glass, although these are very stiff substrates (megapascal, MPa – gigapascal GPa range), and are known to result in non-physiological culture conditions (Jensen and Teng, 2020). Some studies have utilized purely elastic 2D polyacrylamide films coated with various adhesive ECM proteins to investigate HTM cell behaviors (Raghunathan et al., 2013); however, the ECM of native TM tissue is soft (pascal, Pa – kilopascal, kPa range), viscoelastic and 3D. The HTM is composed of specialized HTM cells and ECM proteins such as non-fibrillar and fibrillar collagens, elastic fibrils, proteoglycans and glycosaminoglycans as well as basement membrane-like materials (Abu-Hassan et al., 2014; Acott and Kelley, 2008; Hann and Fautsch, 2011; Keller and Acott, 2013; Tamm, 2009). Viscoelastic 3D hydrogels (i.e., water-swollen networks of polymers) are attractive biomaterials owing to their similarity to *in vivo* cellular microenvironments and tunable physiochemical properties (e.g., stiffness, degradability) (Drury and Mooney, 2003). Thus, tissue-mimetic hydrogels provide an environment that allows for accurate modeling of cell-ECM interactions. We recently reported that a biomimetic HTM hydrogel can be used to investigate 3D cell-ECM interactions under normal and simulated glaucomatous conditions (Li et al., 2021).

Taken together, HTM cells play an essential role in modulating outflow resistance by controlling the production of contractile forces and the secretion/degradation of ECM proteins to support tissue homeostasis (Kelley et al., 2009). TGFβ2-induced HTM cells show increased ECM protein secretion/deposition, actin expression and tension, all strongly associated with HTM cell contractility in POAG (Fleenor et al., 2006; Fuchshofer and Tamm, 2012; Fuchshofer et al., 2003; Han et al., 2011; Pattabiraman and Rao, 2010; Tamm et al., 1999; Tovar-Vidales et al., 2011; Welge-Lüßen et al., 2000). Thus, the purpose of this study was to determine: (a) the molecular mechanism of HTM cell contractility underlying TGFβ2-induced glaucomatous dysfunction and, specifically, whether there is crosstalk between ERK and ROCK signaling pathways, and (b) whether the mechanistic response of HTM cells to TGFβ2 varies depending on the culture substrate (i.e., biomimetic hydrogels composed of ECM biopolymers found in the native tissue in comparison to conventional glass coverslips).

## 2. Materials and Methods

### 2.1 HTM cell isolation and culture

Human donor eye tissue use was approved by the SUNY Upstate Medical University Institutional Review Board (protocol #1211036), and all experiments were performed in accordance with the tenets of the Declaration of Helsinki for the use of human tissue. Primary human trabecular meshwork (HTM) cells were isolated from healthy donor corneal rims discarded after transplant surgery, and cultured according to established protocols (Keller et al., 2018; Stamer et al., 1995). All HTM cell strains used in this study were validated with dexamethasone (DEX; Fisher Scientific, Waltham, MA, USA; 100 nM) induced myocilin (MYOC) expression in more than 50% of cells by immunocytochemistry and immunoblot analyses (**Suppl. Fig. 1** and (Li et al., 2021)). HTM cells were cultured in low-glucose Dulbecco’s Modified Eagle’s Medium (DMEM; Gibco; Thermo Fisher Scientific) containing 10% fetal bovine serum (FBS; Atlanta Biologicals, Flowery Branch, GA, USA) and 1% penicillin/streptomycin/glutamine (PSG; Gibco), and maintained at 37°C in a humidified atmosphere with 5% CO2. Fresh media was supplied every 2-3 days. Three different HTM cell strains were used herein, and all studies were conducted using cells passage 3-6. Donor demographics were as follows: Female/60 years old; Female/50 years old; Male/34 years old.

### 2.2 Preparation of hydrogels

Hydrogel precursors methacrylate-conjugated bovine collagen type I (MA-COL, Advanced BioMatrix, Carlsbad, CA, USA; 3.6 mg/ml [all final concentrations]), thiol-conjugated hyaluronic acid (SH-HA, Glycosil^®^, Advanced BioMatrix; 0.5 mg/ml, 0.025% (w/v) photoinitiator Irgacure^®^ 2959; Sigma-Aldrich, St. Louis, MO, USA) and in-house expressed elastin-like polypeptide (ELP, thiol via KCTS flanks (Li et al., 2021); 2.5 mg/ml) were thoroughly mixed. Thirty microliters of the hydrogel solution were pipetted onto a Surfasil (Fisher Scientific) coated 12-mm round glass coverslip followed by placing a regular 12-mm round glass coverslip onto the hydrogels. Constructs were crosslinked by exposure to UV light (OmniCure S1500 UV Spot Curing System; Excelitas Technologies, Mississauga, Ontario, Canada) at 320-500 nm, 2.2 W/cm^2^ for 5 s, as previously described (Li et al., 2021). The hydrogel-adhered coverslips were removed with fine-tipped tweezers and placed in 24-well culture plates (Corning; Thermo Fisher Scientific). HTM cell-laden hydrogels were prepared by mixing HTM cells (1.0 × 10^6^ cells/ml) with MA-COL (3.6 mg/ml [all final concentrations]), SH-HA (0.5 mg/ml, 0.025% (w/v) photoinitiator) and ELP (2.5 mg/ml) on ice, followed by pipetting 10 μl droplets of the HTM cell-laden hydrogel precursor solution onto polydimethylsiloxane-coated (Sylgard 184; Dow Corning) 24-well culture plates (**Suppl. Fig. 2**).

### 2.3 HTM cell treatments

HTM cells were seeded at 2 × 10^4^ cells/cm^2^ on premade hydrogels/coverslips/tissue culture plates (**Suppl. Fig. 2**), and cultured in DMEM with 10% FBS and 1% PSG for 1 or 2 days to grow to 80 – 90 % confluence. Then, HTM cells were cultured in serum-free DMEM with 1% PSG and subjected to the following treatments for 2 h (starvation in serum-free DMEM with 1% PSG for 24 h prior to treatments) or 3 d: 1) vehicle control, 2) TGFβ2 (2.5 ng/ml; R&D Systems, Minneapolis, MN, USA), 3) TGFβ2 (2.5 ng/ml) + U0126 (10 μM; Promega, Madison, WI, USA), 4) TGFβ2 (2.5 ng/ml) + Y27632 (10 μM; Sigma-Aldrich). The monolayer HTM cells were processed for immunoblot and immunocytochemistry analyses.

### 2.4 Immunoblot analysis

Protein was extracted from cells grown on 6-well culture plates using lysis buffer (CelLytic^™^ M, Sigma-Aldrich) supplemented with Halt™ protease/phosphatase inhibitor cocktail (Thermo Fisher Scientific). Equal protein amounts (10 μg), determined by standard bicinchoninic acid assay (Pierce; Thermo Fisher Scientific), in 4× loading buffer (Invitrogen; Thermo Fisher Scientific) with 5% beta-mercaptoethanol (Fisher Scientific), were boiled for 5 min and subjected to SDS-PAGE using NuPAGE™ 4-12% Bis-Tris Gels (Invitrogen; Thermo Fisher Scientific) at 120V for 80 min and transferred to 0.45 μm PVDF membranes (Sigma; Thermo Fisher Scientific). Membranes were blocked with 5% bovine serum albumin (Thermo Fisher Scientific) in trisbuffered saline with 0.2% Tween^®^20 (Thermo Fisher Scientific), and probed with primary antibodies against p-ERK (anti-p-ERK [9101S] 1:4000; Cell Signaling Technology, Danvers, MA, USA), ERK (anti-ERK [9102S] 1:3000; Cell Signaling), and MYOC (anti-MYOC [MABN866] 1:2000; Sigma-Aldrich) followed by incubation with an HRP-conjugated secondary antibody (Cell Signaling). Bound antibodies were visualized with the enhanced chemiluminescent detection system (Pierce) on autoradiography film (Thermo Fisher Scientific). Densitometry of scanned films was performed using Fiji software (NIH). Data were normalized to GAPDH (anti-GAPDH [G9545] 1:40000; Sigma-Aldrich).

### 2.5 Immunocytochemistry analysis

HTM cells in presence of the different treatments were fixed with 4% paraformaldehyde (Thermo Fisher Scientific) at room temperature for 20 min, permeabilized with 0.5% Triton™ X-100 (Thermo Fisher Scientific), blocked with blocking buffer (BioGeneX), and incubated with primary antibodies against MYOC (anti-MYOC [ab41552] 1:200; Abcam), α-smooth muscle actin (anti-αSMA [C6198] 1:500; Sigma-Aldrich), fibronectin (FN [ab45688] 1:500; Abcam), or phospho-myosin light chain 2 (Ser19) (p-MLC [3675] 1:200; Cell Signaling), followed by incubation with Alexa Fluor^®^ 488-conjugated or 594-conjugated secondary antibodies (Invitrogen; Thermo Fisher Scientific); nuclei were counterstained with 4’,6’-diamidino-2-phenylindole (DAPI; Abcam). Similarly, cells were stained with Phalloidin-iFluor 488 (Abcam)/DAPI according to the manufacturer’s instructions. Coverslips were mounted with ProLong™ Gold Antifade (Invitrogen) on Superfrost™ microscope slides (Fisher Scientific), and fluorescent images were acquired with an Eclipse N*i* microscope (Nikon Instruments, Melville, NY, USA). Fluorescent signal intensity was measured using Fiji software (National Institutes of Health (NIH), Bethesda, MD, USA). Quantitative immunostaining was measured for F-actin, αSMA, FN, and p-MLC from 10 images/group following image background subtraction. F-actin fiber alignment was measured using the Directionality Fourier transform (FFT) plug-in in Fiji. The nuclear shape index (circularity = 4*π*area/perimeter^2^) and nuclear aspect ratio (major axis/minor axis), indicators of the shape of the cells, were also measured using Fiji software.

### 2.6 HTM hydrogel contraction analysis

HTM cell-laden hydrogels were cultured in DMEM with 10% FBS and 1% PSG in presence of the different treatments. Longitudinal brightfield images were acquired at 0 d and 7 d with an Eclipse T*i* microscope (Nikon). Construct area was measured using Fiji software (NIH) and normalized to 0 d followed by normalization to controls.

### 2.7 HTM hydrogel cell proliferation analysis

Cell proliferation was measured with the CellTiter 96^®^ Aqueous Non-Radioactive Cell Proliferation Assay (Promega) following the manufacturer’s protocol. HTM hydrogels cultured in DMEM with 10% FBS and 1% PSG in presence of the different treatments for 7 d (N = 3 per group) were incubated with the staining solution (38 μl MTS, 2 μl PMS solution, 200 μl DMEM) at 37°C for 1.5 h. Absorbance at 490 nm was recorded using a spectrophotometer plate reader (BioTEK, Winooski, VT, USA). Blank-subtracted absorbance values served as a direct measure of HTM cell proliferation.

### 2.8 Statistical analysis

Individual sample sizes are specified in each figure caption. Comparisons between groups were assessed by one-way analysis of variance with Tukey’s multiple comparisons *post hoc* tests, as appropriate. All data are shown with mean ± SD, some with individual data points. The significance level was set at p<0.05 or lower. GraphPad Prism software v9.1 (GraphPad Software, La Jolla, CA, USA) was used for all analyses.

## 3. Results

### 3.1 Effects of TGFβ2 in absence or presence of ERK or ROCK inhibition on F-actin stress fibers in HTM cells

To investigate the effects of different cell culture substrates and downstream signaling events of TGFβ2 stimulation, HTM cells were seeded on top of pre-made hydrogels or glass coverslips (**Suppl. Fig. 2**), and treated with TGFβ2 alone, or co-treated with ERK inhibitor U0126 or ROCK inhibitor Y27632 and then stained for actin structures. We observed significantly more F-actin stress fibers in HTM cells on hydrogels treated with TGFβ2 vs. controls, while co-treatment with U0126 or Y27632 restored F-actin intensity to control levels (**Fig. 1A,B**). Likewise, for HTM cells plated on glass, we observed that TGFβ2 significantly increased F-actin compared to controls. We did not detect a difference between cells co-treated with TGFβ2 + U0126 vs. TGFβ2-treated cells. However, Y27632 treatment partially prevented changes induced by TGFβ2, albeit not to control levels (**Fig. 1C**).

**Fig. 1.**
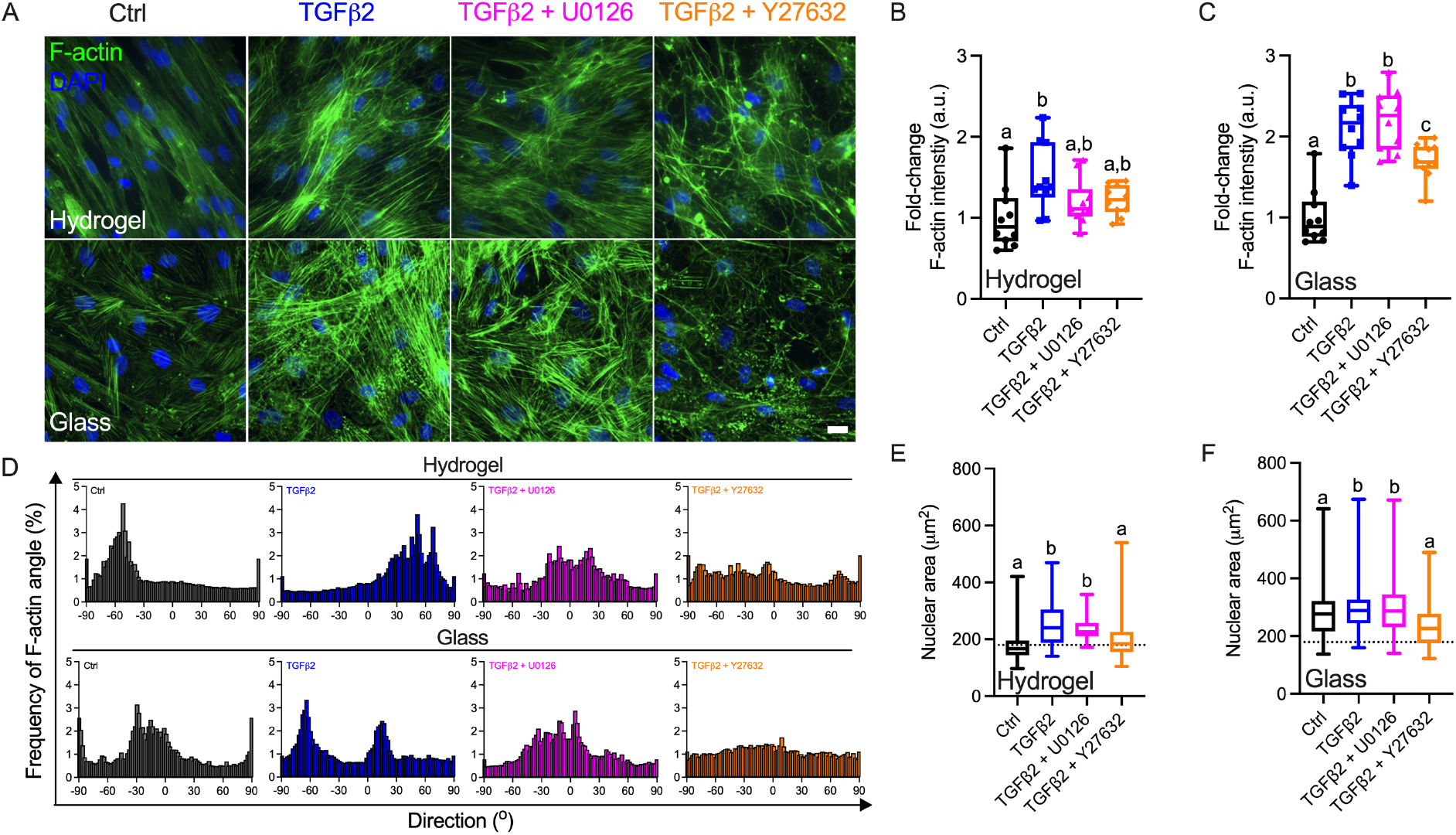
Effects of TGFβ2 in absence or presence of ERK or ROCK inhibition on F-actin stress fibers and morphology of HTM cells. **(A)** Representative fluorescence micrographs of F-actin in HTM cells on hydrogel and glass substrates subjected to control, TGFβ2 (2.5 ng/mL), TGFβ2 + U0126 (10 μM), TGFβ2 + Y27632 (10 μM) at 3 d (F-actin = green; DAPI = blue). Scale bar, 20 μm. Quantification of F-actin intensity in (**B**) HTM cells on hydrogels and (**C**) HTM cells on glass subjected to the different treatments at 3 d (n = 10 images from three biological replicates). **(D)** Histograms representing the F-actin direction in HTM cells on hydrogel and glass substrates subjected to the different treatments. A unimodal distribution denotes aligned uniaxial F-actin fiber alignment, whereas a multimodal distribution indicates anisotropic F-actin array arrangement. Quantification of nuclear area in (**E**) HTM cells on hydrogels and (**F**) HTM cells on glass subjected to the different treatments at 3 d (n = 100 nuclei from three biological replicates; dotted lines show mean value of control HTM cells on hydrogel for reference). In B, C, E and F, the box and whisker plots represent median values (horizontal bars), 25th to 75th percentiles (box edges) and minimum to maximum values (whiskers), with all points plotted in B, C; shared significance indicator letters represent non-significant difference (p>0.05), distinct letters represent significant difference (p<0.05).

We used the FIJI plugin “Directionality Fourier transform (FFT)” to quantify the relative orientation of actin filaments in HTM cells. In general terms, this parameter represents perfectly aligned F-actin (i.e., parallel) as a histogram with a single unimodal peak and anisotropic filaments as deviations i.e., less aligned, multimodal peaks (Deravi et al., 2017; Wang et al., 2014). In our analysis, HTM cells on hydrogels (Control) displayed highly aligned F-actin with a unimodal histogram, whereas F-actin in HTM cells treated with TGFβ2 ± U0126/Y27632 had misordered F-actin arrays, represented as multimodal peaks (**Fig. 1A**). Additionally, all cells plated on glass regardless of treatment displayed anisotropic F-actin arrays (**Fig. 1D**).

Next, we assessed effects of the different substrates (i.e., hydrogel and glass) and treatments on HTM cell morphology. HTM cells on hydrogels with TGFβ2 ± U0126 treatment exhibited a significantly increased nuclear area compared to controls (**Fig. 1E**) with an increased nuclear shape index (NSI) and a reduced nuclear aspect ratio (**Suppl. Fig. 3A,B**). By contrast, Y27632 significantly prevented changes induced by TGFβ2. HTM cells on glass across groups displayed overall larger nuclear area and higher NSI, indicative of bigger and more rounded nuclei vs. cells on hydrogels (indicated with the dashed line in **Fig. 1E,F** and **Suppl. Fig. 3A-D**). Mirroring the behavior on hydrogels, treatment with TGFβ2 ± U0126 significantly increased HTM cell nuclear area vs. controls on glass, whereas Y27632 treatment restored nuclear size to control levels. HTM cells on glass subjected to the different treatments showed no significant difference in NSI, and only cells treated with TGFβ2 + U0126 exhibited significantly lower nuclear aspect ratio compared to TGFβ2 + Y27632 treated cells. (**Fig. 1E,F; Suppl. Fig. 3**).

Together, these data show that HTM cells on biomimetic soft ECM substrates in absence of treatment exhibit a highly aligned actin cytoskeleton, while cells on supraphysiologic stiff substrates display differential F-actin organization. Exposure to the different treatments results in a generally lesser organized cytoskeleton across groups, suggesting that the biochemical cues impart cellular changes that supersede effects mediated by the culture substrates. TGFβ2 reliably increases actin stress fibers, which is reduced by Y27632, while ERK inhibition has little-to-no effect. Nevertheless, HTM cells display distinct nuclear size/shape in soft vs. stiff culture environments, suggesting that overall cell morphology is partly affected by the underlying substrate.

### 3.2 Effects of TGFβ2 in absence or presence of ERK or ROCK inhibition on αSMA expression in HTM cells

To determine the crosstalk between ERK and ROCK signaling pathways upon TGFβ2 stimulation, HTM cells were treated with TGFβ2 in the presence of U0126 or Y27632 on tissue culture plastic to assess ERK and ROCK protein expression. We observed that TGFβ2 significantly increased ERK1/2 phosphorylation (=activation) vs. controls. As expected, cotreatment with U0126 prevented ERK activation, while Y27632 did not affect the TGFβ2-induced phosphorylation of ERK (**Fig. 2A,B**).

**Fig. 2.**
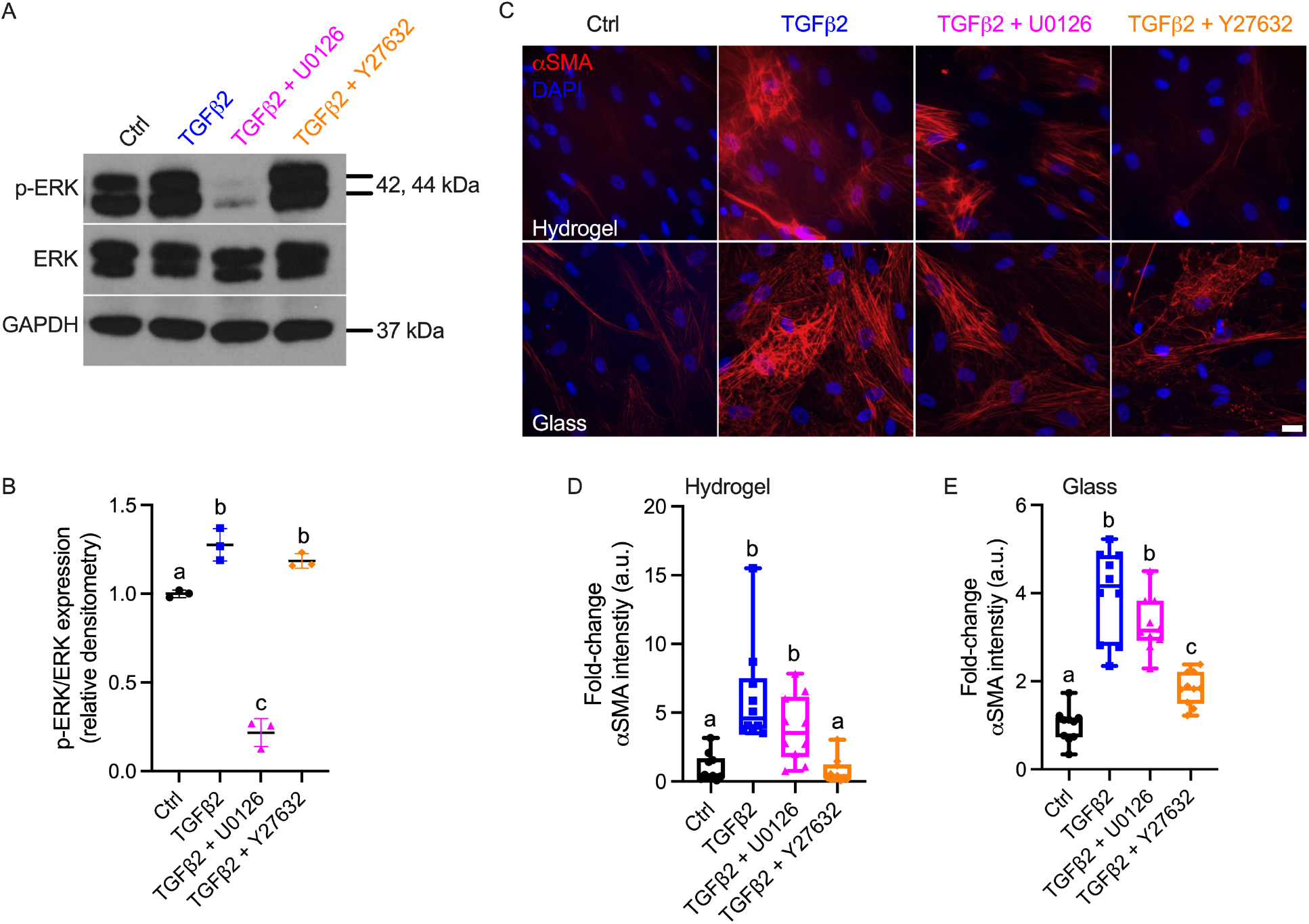
Effects of TGFβ2 in absence or presence of ERK or ROCK inhibition on αSMA expression in HTM cells. (A) Immunoblot of p-ERK and ERK. (B) Quantification of p-ERK/ERK (n = 3 per group). **(C)** Representative fluorescence micrographs of αSMA in HTM cells on hydrogel and glass substrates subjected to control, TGFβ2 (2.5 ng/ml), TGFβ2 + U0126 (10 μM), TGFβ2 + Y27632 (10 μM) at 3 d (αSMA = red; DAPI = blue). Scale bar, 20 μm. Quantification of αSMA intensity in (**D**) HTM cells on hydrogels and (**E**) HTM cells on glass subjected to the different treatments at 3 d (n = 10 images from three biological replicates). The box and whisker plots represent median values (horizontal bars), 25th to 75th percentiles (box edges) and minimum to maximum values (whiskers), with all points plotted; shared significance indicator letters represent non-significant difference (p>0.05), distinct letters represent significant difference (p<0.05).

TGFβ2 has been shown to induce HTM cell trans-differentiation, as indicated by expression of the myofibroblast marker α-smooth muscle actin (αSMA) (Pattabiraman and Rao, 2010; Sun et al., 2016; Zhao et al., 2004). To gain insight into the effects of ERK and ROCK inhibition on HTM cells, we next assessed αSMA levels in HTM cells treated with TGFβ2 ± U0126 or Y27632. TGFβ2 ± U0126 treated groups showed significantly increased αSMA expression in HTM cells vs. the other groups independent of the culture substrate (**Fig. 2C-E**). We observed that TGFβ2 induced a bigger change of αSMA expression in HTM cells on hydrogels (6.15-fold of controls) compared to glass (3.99-fold of controls). Y27632 restored TGFβ2-induced αSMA expression to control levels in HTM cells on hydrogels (**Fig. 2C,D**). Likewise, Y27632 treatment significantly decreased αSMA expression vs. the TGFβ2-treated group in HTM cells on glass, but did not fully reach baseline levels (**Fig. 2C,E**).

Collectively, these data show that TGFβ2 upregulates ERK activity in HTM cells, which is potently attenuated by U0126, while ROCK inhibition has no effect on TGFβ2-induced ERK activation. Furthermore, TGFβ2 increases αSMA expression, which is prevented by Y27632, whereas ERK inhibition does not affect aberrant αSMA stress fiber formation independent of the underlying substrate. This suggests that αSMA expression is largely unaffected by the biophysical cues from the culture environment.

### 3.3 Effects of TGFβ2 in absence or presence of ERK or ROCK inhibition on FN synthesis in HTM cells

There is increased accumulation of ECM proteins in the HTM tissue of glaucomatous eyes, among which fibronectin (FN) is a major component (Hann et al., 2001; Stamer and Clark, 2017). To assess the effect of TGFβ2 on ECM protein expression and deposition, we evaluated FN levels in HTM cells subjected to the different treatments. We observed overall more pronounced FN fibril formation in HTM cells grown on glass vs. hydrogels in absence of any treatment, suggesting that ECM deposition was affected by the underlying substrate (**Fig. 3A**). Importantly, TGFβ2-treated HTM cells showed significantly increased FN expression and deposition vs. controls independent of the culture substrate, whereas U0126 and Y27632 markedly reduced the FN signal intensity, indicating that ERK and ROCK activities were associated with ECM expression and deposition (**Fig. 3**).

**Fig. 3.**
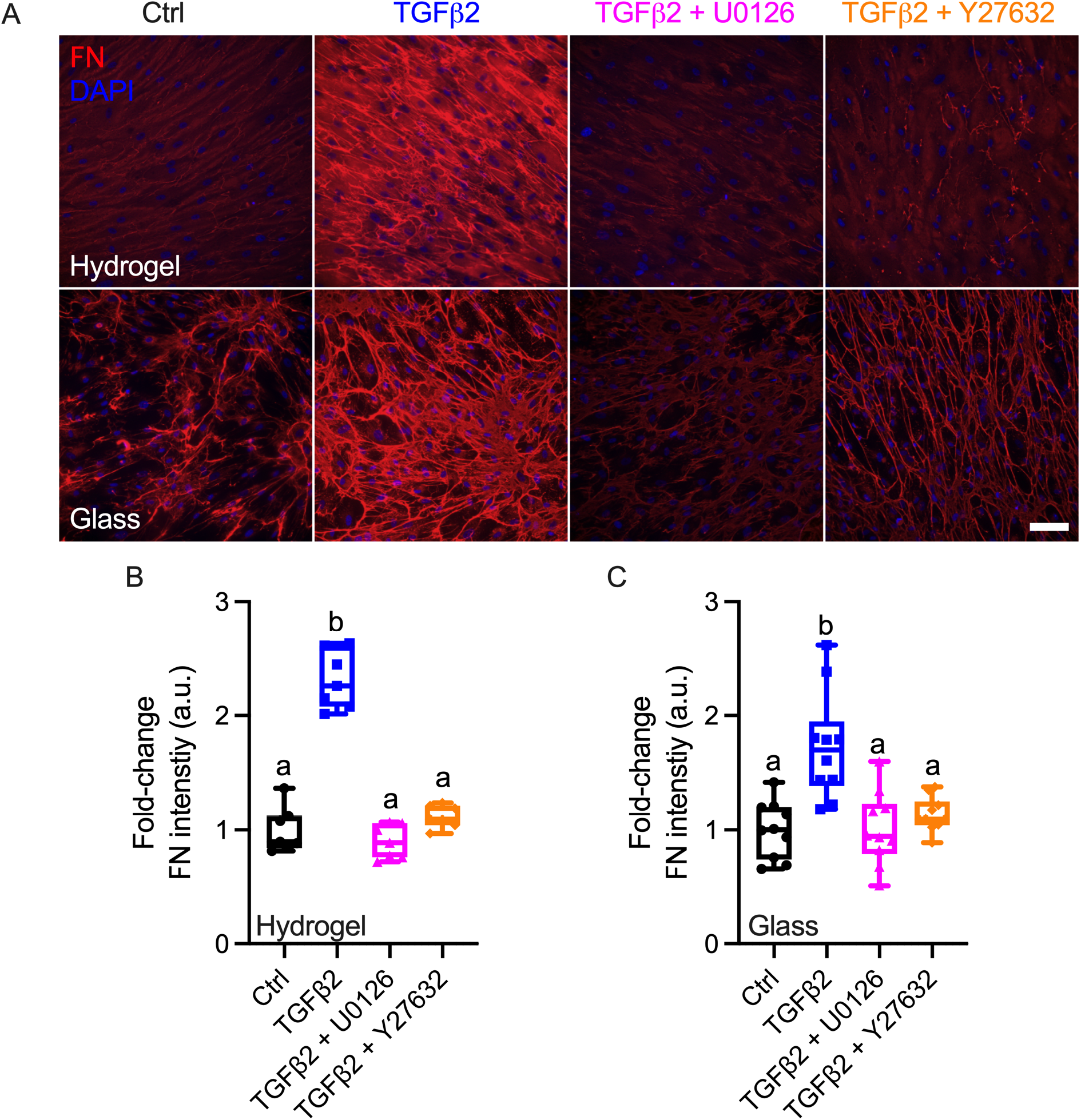
Effects of TGFβ2 in absence or presence of ERK or ROCK inhibition on FN synthesis in HTM cells. **(A)** Representative fluorescence micrographs of FN in HTM cells on hydrogel and glass substrates subjected to control, TGFβ2 (2.5 ng/ml), TGFβ2 + U0126 (10 μM), TGFβ2 + Y27632 (10 μM) at 3 d (FN = red; DAPI = blue). Scale bar, 100 μm. Quantification of FN intensity in (**B**) HTM cells on hydrogels and (**C**) HTM cells on glass subjected to the different treatments at 3 d (n = 10 images from three biological replicates). The box and whisker plots represent median values (horizontal bars), 25th to 75th percentiles (box edges) and minimum to maximum values (whiskers), with all points plotted; shared significance indicator letters represent non-significant difference (p>0.05), distinct letters represent significant difference (p<0.05).

Together, these data show that TGFβ2 increases FN expression/deposition in HTM cells, which is potently attenuated by either U0126 or Y27632 co-treatment independent of the underlying culture substrate. This suggests that ERK and ROCK activation are prerequisite for TGFβ2-induced aberrant FN fibril formation in HTM cells.

### 3.4 Effects of TGFβ2 in absence or presence of ERK or ROCK inhibition on p-MLC expression in HTM cells

The Rho/ROCK pathway is a master regulator of the actin cytoskeleton and cell contractility via phosphorylation of myosin light chain (MLC), one of its main substrates (Wang et al., 2013). ROCK, which has two isoforms (ROCK1 and ROCK2) (Liu et al., 2018), is strongly associated with HTM cell contractility; we have demonstrated that TGFβ2 increases HTM cell contractility, which is potently rescued by ROCK inhibition (Li et al., 2021). To investigate the effects of ERK and ROCK inhibition on HTM cell contractility, ROCK expression levels in HTM cells treated with TGFβ2 in the presence or absence of U0126 or Y27632 were assessed. TGFβ2-treated samples showed qualitatively more ROCK1 expression vs. control (~1.3-fold), which was potentiated with U0126 co-treatment (~1.5-fold of controls) while Y27632 did not influence ROCK1 expression; ROCK2 levels were comparable across all groups (**Suppl. Fig. 4**).

Next, we assessed the expression of p-MLC. TGFβ2 significantly increased p-MLC levels vs. controls in HTM cells on both hydrogels and glass, which was completely prevented by Y27632 co-treatment. Strikingly, HTM cells on hydrogels co-treated with TGFβ2 and U0126 exhibited significantly increased expression of p-MLC vs. TGFβ2-treated samples (**Fig. 4A,B**), while U0126 did not affect TGFβ2-induced p-MLC expression in HTM cells on glass (**Fig. 4B,D**).

**Fig. 4.**
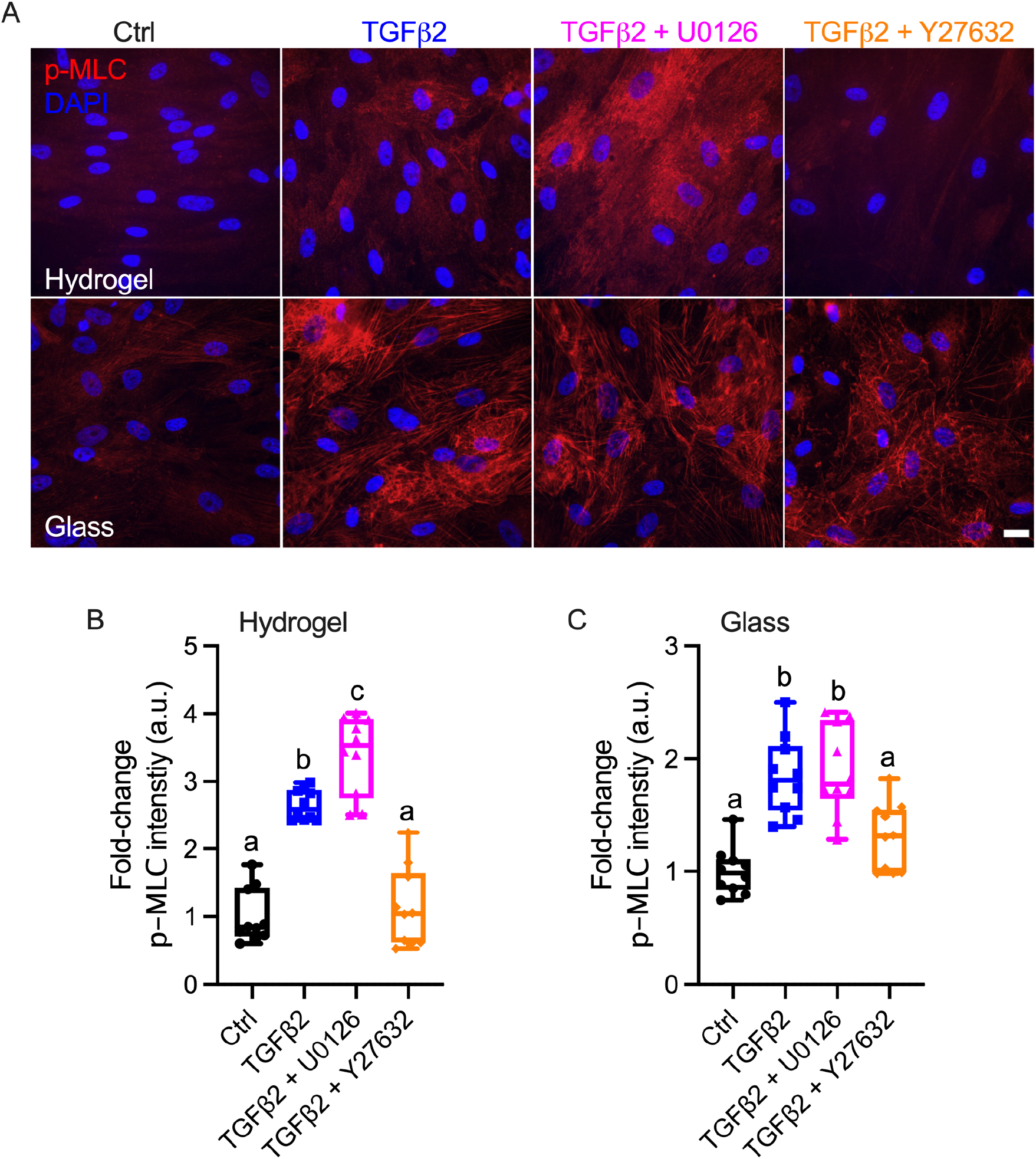
Effects of TGFβ2 in absence or presence of ERK or ROCK inhibition on p-MLC expression in HTM cells. **(A)** Representative fluorescence micrographs of p-MLC in HTM cells on hydrogel and glass substrates subjected to control, TGFβ2 (2.5 ng/ml), TGFβ2 + U0126 (10 μM), TGFβ2 + Y27632 (10 μM) at 3 d (p-MLC = red; DAPI = blue). Scale bar, 20 μm. Quantification of p-MLC intensity in (**B**) HTM cells on hydrogels and (**C**) HTM cells on glass subjected to the different treatments at 3 d (n = 10 images from three biological replicates). The box and whisker plots represent median values (horizontal bars), 25th to 75th percentiles (box edges) and minimum to maximum values (whiskers), with all points plotted; shared significance indicator letters represent non-significant difference (p>0.05), distinct letters represent significant difference (p<0.05).

Collectively, these data show that TGFβ2 elevates p-MLC levels via ROCK1, which is further potentiated with U0126 co-treatment in HTM cells on biomimetic soft ECM substrates but not on supraphysiologic stiff substrates. This suggests that active ERK signaling is required to prevent aberrant ROCK activation that exacerbates TGFβ2-induced HTM cell contractility.

### 3.5 Effects of TGFβ2 in absence or presence of ERK or ROCK inhibition on HTM hydrogel contractility

In our recent study, we showed that HTM cells encapsulated in ECM biopolymer hydrogels are highly contractile, and that TGFβ2 increases 3D HTM hydrogel contraction vs. controls (Li et al., 2021). To investigate the effects of ERK and ROCK signaling, and their crosstalk, on HTM cell contractility, we assessed HTM hydrogel contraction upon treatment with TGFβ2 in the presence or absence of either U0126 or Y27632. TGFβ2-treated HTM hydrogels exhibited significantly greater contraction vs. controls by 7 d (77.29%), whereas Y27632 significantly decreased hydrogel contraction (=relaxed) in absence (117.90%) or presence (114.10%) of TGFβ2 stimulation, consistent with our previous report. Interestingly, U0126 treatment alone induced significantly greater contraction (65.91%) compared with TGFβ2-treated samples, and cotreatment of TGFβ2 + U0126 induced even higher HTM hydrogel contraction (49.68%). Hydrogel size with TGFβ2 + U0126 + Y27632 treatment was similar to controls (**Fig. 5A, B**), indicating that hydrogel relaxation via ROCK inhibition can overcome HTM cell hypercontractility mediated by ERK inhibition.

**Fig. 5.**
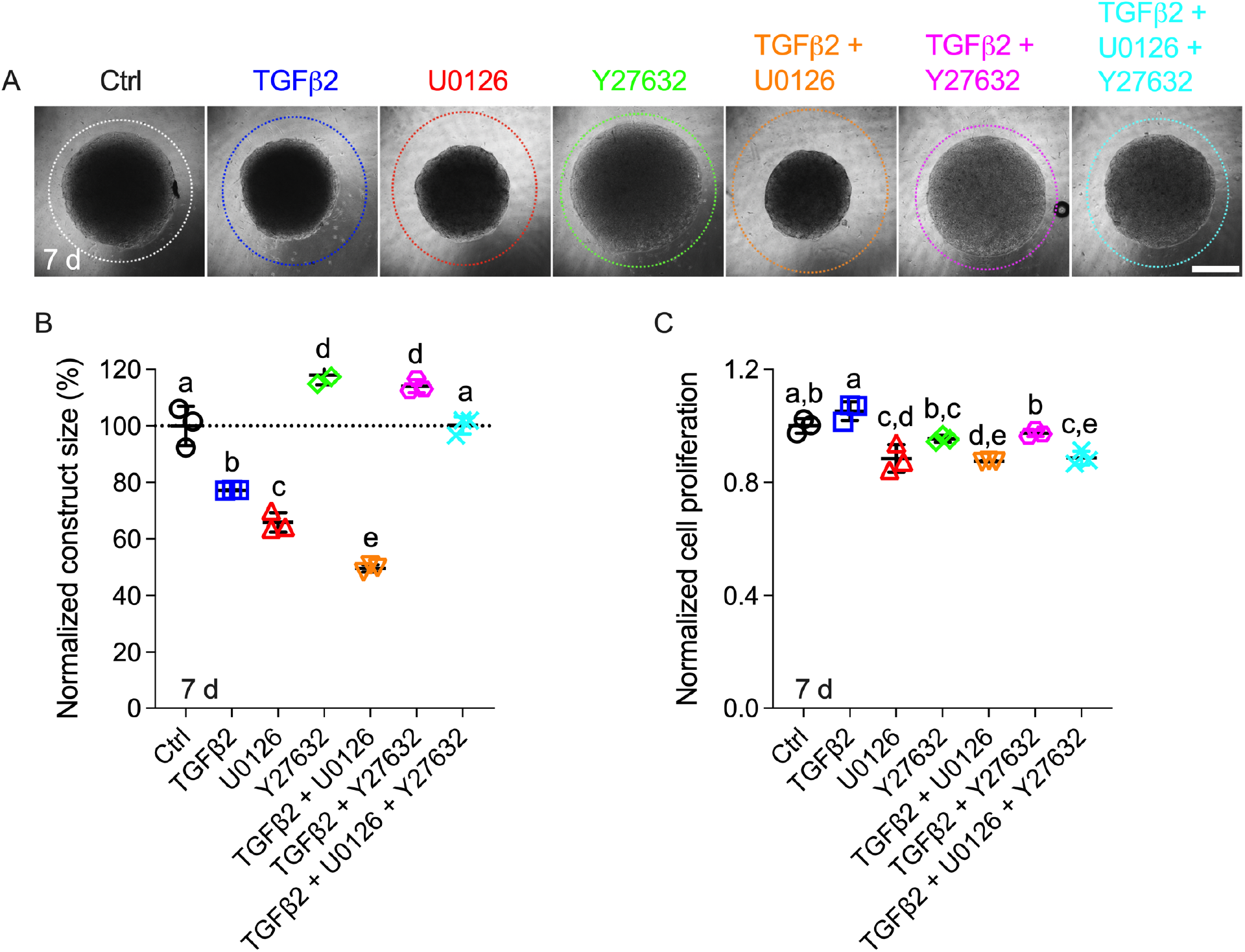
Effects of TGFβ2 in absence or presence of ERK or ROCK inhibition on HTM hydrogel contractility. **(A)** Representative brightfield images of HTM hydrogels subjected to control, TGFβ2 (2.5 ng/ml), TGFβ2 + U0126 (10 μM), TGFβ2 + Y27632 (10 μM) at 7 d (dashed lines outline original size of constructs at 0 d. Scale bar, 1 mm). **(B)** Construct size quantification of HTM hydrogels subjected to the different treatments at 7 d (n = 3 per group; dotted line shows control value for reference; shared significance indicator letters represent non-significant difference (p>0.05), distinct letters represent significant difference (p<0.05)). **(C)** Cell proliferation quantification of HTM hydrogels subjected to the different treatments at 7 d (n = 3 per group; shared significance indicator letters represent non-significant difference (p>0.05), distinct letters represent significant difference (p<0.05)).

To determine if hydrogel contractility was influenced by the cell number, we assessed HTM cell proliferation in constructs subjected to the different treatments. HTM cells in hydrogels treated with TGFβ2, TGFβ2 + U0126 or TGFβ2 + U0126 + Y27632 showed significantly less proliferation compared to controls at 7 d, in spite of these constructs exhibiting greater contraction vs. controls (**Fig. 5C**).

Together, these data show that TGFβ2 robustly induces HTM hydrogel contractility, which is exacerbated by U0126, while ROCK inhibition potently relaxes the HTM hydrogels. This suggests that active ERK signaling is necessary to prevent abnormal ROCK activation that drives pathologic TGFβ2-induced HTM cell hypercontractility in a tissue-mimetic 3D environment, consistent with the p-MLC expression data.

## 4. Discussion

TGFβ2 is a major contributor to the pathologic changes occurring in HTM cells/tissue in POAG (Fuchshofer and Tamm, 2009; Quigley, 1993). A number of studies have shown elevated levels of TGFβ2 in the aqueous humor of glaucomatous patients compared to age-matched normal eyes (Agarwal et al., 2015; Inatani et al., 2001; Ochiai and Ochiai, 2002; Picht et al., 2001). TGFβ2 induces the formation of F-actin stress fibers and expression of αSMA, as well as increases the synthesis and decreases the degradation of several ECM proteins, including FN, produced by HTM cells - all strongly associated with pathologic cell contractility and outflow resistance (Alexander et al., 1998; Fleenor et al., 2006; Han et al., 2011; Pattabiraman and Rao, 2010). In our recent study, we demonstrated increased contraction of HTM cell-laden hydrogels in response to glaucomatous stressors dexamethasone and TGFβ2 (Li et al., 2021). Here, our aim was to elucidate mechanisms governing the HTM cell contractility response to simulated glaucomatous conditions, with a focus on TGFβ2 induction.

TGFβ2 is known to signal via both canonical Smad and non-canonical signaling pathways, such as ERK and ROCK; both have been shown to regulate F-actin, αSMA and FN expression (Zhang, 2009). It has also been reported that ROCK promotes smooth muscle cell migration and serotonin-mediated cell proliferation via upregulating ERK activity (Liu et al., 2004; Zhao et al., 2012). In neurons and microglia, ERK activity is negatively regulated by ROCK (Hensel et al., 2015). Thus, there is mounting evidence that ERK and ROCK signaling pathways crosstalk to regulate behaviors in a variety of cells (Hensel et al., 2015; Liu et al., 2004; Tong et al., 2016; Zhao et al., 2012), whereas mechanisms of their crosstalk in HTM cells are less well understood. TGFβ2 has been shown to upregulate ERK via activating RhoA/ROCK in HTM cells to increase ECM protein expression, F-actin stress fibers and αSMA (Pattabiraman and Rao, 2010), putting the two non-canonical signaling pathways in sequence. By contrast, we here demonstrate that ROCK inhibition via Y27632 did not influence TGFβ2-induced ERK activity, as determined by measuring levels of p-ERK (**Fig. 2A,B**).

Most *in vitro* studies of HTM cell (patho-)physiology have relied on conventional 2D cell monolayer cultures on plastic or glass. However, biophysical cues such as substrate composition and stiffness are known to be potent modulators of cell behaviors (Raghunathan et al., 2013). ECM protein-based hydrogels are widely used to mimic the natural tissue environment and allow for more accurate investigation of cellular behaviors *in vitro* (Green and Elisseeff, 2016; Khademhosseini and Langer, 2007; Lee and Mooney, 2001; Seliktar, 2012; Zhang and Khademhosseini, 2017). Importantly, these types of hydrogels allow for seeding cells atop and encapsulating within the 3D polymer network to establish artificial tissue models without further surface modifications, unlike polyacrylamide gels which can only be used in 2D cell culture and need ECM protein modification. In our recent study, we established a bioengineered HTM hydrogel formed by mixing normal donor-derived HTM cells with collagen type I, elastin-like polypeptide and hyaluronic acid, each containing photoactive functional groups, followed by UV crosslinking. Our *in vitro* HTM model can be used to investigate 3D cell-ECM interactions under normal and simulated glaucomatous conditions (Li et al., 2021).

We first compared the behaviors of HTM cells cultured atop the biomimetic hydrogels with cells on traditional glass coverslips. HTM cells on hydrogels showed aligned F-actin fibers, while HTM cells on glass showed multiple directions of F-actin fibers in absence of exogenous biochemical cues (**Fig. 1A,D**). Besides, HTM cells on glass showed bigger and increased spherical nuclei compared with HTM cells on hydrogels, which demonstrated that HTM cells exhibited distinct morphology on different substrates (**Fig. 1E,F; Suppl. Fig. 3**). In general, TGFβ2 increased F-actin and αSMA expression, which was largely prevented by Y27632, while ERK inhibition had no effect on TGFβ2-induced actin cytoskeletal changes, suggesting that F-actin and αSMA expression is largely unaffected by the underlying substrate (**Fig. 1 A,C,D and Fig. 2C-E**).

ECM remodeling is critical for HTM function, and increased FN expression/deposition has been associated with glaucomatous tissue dysfunction (Stamer and Clark, 2017). We showed higher FN deposition in HTM cells on glass compared to hydrogels (**Fig. 3**). It is possible that the artificial culture conditions of glass coverslips (i.e., 2D, supraphysiological stiffness) cause higher cell stress in HTM cells potentially instructing them to make their own softer substrate, which leads to more FN fibril formation vs. HTM cells grown on biomimetic soft ECM. FN has multiple isoforms generated by alternative processing of a single primary transcript at three domains: extra domain A (EDA), extra domain B (EDB), and the type III homologies connecting segment (Serini et al., 1998). It has been demonstrated that FN-EDA was elevated in glaucomatous vs. normal HTM tissue (Medina-Ortiz et al., 2013). Future studies will investigate the effects of TGFβ2 in absence or presence of ERK/ROCK inhibition on FN-EDA and FN-EDB.

Phospho-MLC, a downstream target of ROCK activation, induces the generation of cellular forces from actomyosin filament contraction (Lampi and Reinhart-King, 2018; Wang et al., 2013); our results showed that ERK inhibition further increased TGFβ2-induced p-MLC levels in HTM cells on hydrogels, which was not observed in HTM cells on glass (**Fig. 4**). To our knowledge, this is the first report demonstrating that ERK inhibition can increase ROCK and p-MLC levels in HTM cells. In a previous study, it has been shown that both ERK and ROCK inhibition downregulated p-MLC expression in HTM cells cultured on conventional stiff 2D substrates (Pattabiraman and Rao, 2010). Here, we showed that ERK inhibition had no effect on TGFβ2-induced p-MLC in HTM cells on glass (**Fig. 4A,C**), suggesting that p-MLC expression is highly influenced by the underlying substrate. Consistent with the p-MLC data, we showed further increased 3D HTM hydrogel contraction with ERK inhibition, whereas ROCK inhibition had precisely the opposite effects and potently relaxed the TGFβ2-induced hydrogels (**Fig. 5**). Taken together, this suggests that ERK signaling negatively regulates ROCK-mediated (i.e., via p-MLC) HTM cell contractility, and that impairment of this crosstalk balance can contribute to pathologic contraction in the context of the glaucomatous stressor TGFβ2.

In summary, we showed that different culture substrates affected HTM cell morphology and F-actin organization under normal and simulated POAG conditions using TGFβ2 induction. We demonstrated that TGFβ2 increased HTM cell contractility via ERK and ROCK signaling pathways by regulating F-actin, αSMA, FN and p-MLC. Importantly, we showed that ROCK activity was negatively regulated by ERK with TGFβ2 induction (**Fig. 6**). These findings emphasize the critical importance of using 3D viscoelastic ECM substrates for investigating HTM cell physiology and glaucomatous pathophysiology *in vitro*.

**Fig. 6.**
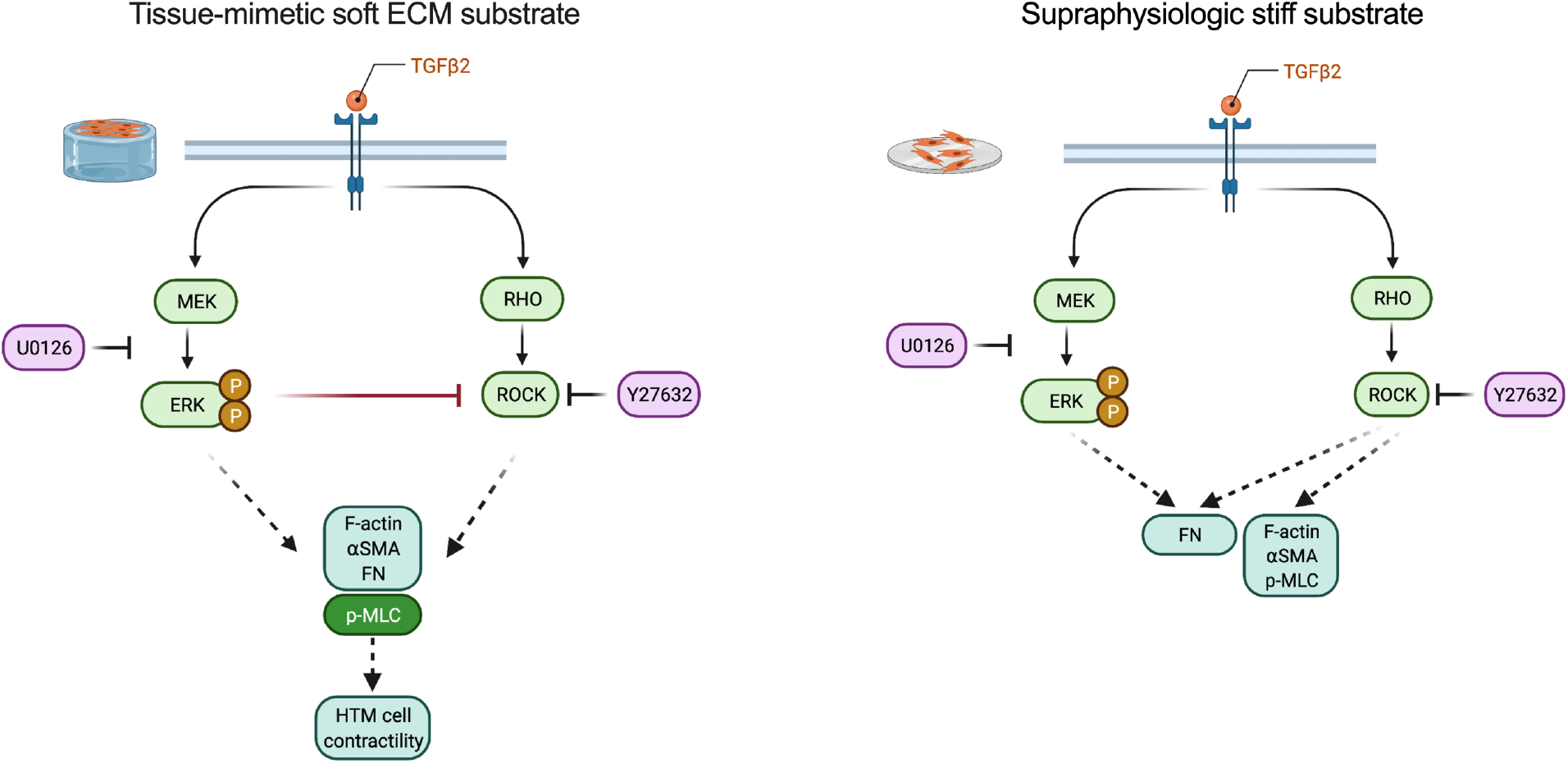
Schematic illustration of how TGFβ2 may regulate HTM cell contractility via ERK and ROCK signaling pathways. HTM cells on hydrogels exhibit aligned F-actin fibers in absence of biochemical cues. Results showed that exposure to TGFβ2 activates ERK and ROCK signaling pathways to regulate F-actin, αSMA, FN and p-MLC, which are associated with pathologic HTM cell contractility under glaucomatous conditions. Importantly, in this tissue-mimetic viscoelastic ECM environment, ROCK activity is negatively regulated by ERK which was not observed in HTM cells on glass, and impairment of this crosstalk balance may contribute to pathologic HTM cell contraction via elevated p-MLC in the context of the known glaucomatous stressor TGFβ2. These findings emphasize the critical importance of using 3D tissue-mimetic soft ECM substrates for investigating HTM cell physiology and glaucomatous pathophysiology *in vitro*. Created with BioRender.com.

## Disclosure

The authors report no conflicts of interest.

## Funding

This project was supported in part by a National Institutes of Health grant GM133485 (to J.L.H-R), an American Glaucoma Society Young Clinician Scientist Award (to P.S.G.), a Syracuse University BioInspired Pilot Grant (to S.H.), unrestricted grants to SUNY Upstate Medical University Department of Ophthalmology and Visual Sciences from Research to Prevent Blindness (RPB) and from Lions Region 20-Y1, and RPB Career Development Awards (to P.S.G. and S.H.).

## Acknowledgments

We thank Dr. Robert W. Weisenthal and the team at Specialty Surgery Center of Central New York for assistance with corneal rim specimens. We also thank Dr. Nasim Annabi at the University of California – Los Angeles for providing the KCTS-ELP, and Drs. Audrey M. Bernstein and Mariano S. Viapiano at Upstate Medical University for imaging support. Lastly, we thank Dr. W. Daniel Stamer at Duke University for the helpful discussions and editing of the final manuscript.

## Author contributions

H.L., J.L.H-R., P.S.G., and S.H designed all experiments, collected, analyzed, and interpreted the data. H.L. and S.H. wrote the manuscript. All authors commented on and approved the final manuscript. P.S.G. and S.H. conceived and supervised the research.

## Competing interests

The authors declare no conflict of interest.

## Data and materials availability

All data needed to evaluate the conclusions in the paper are present in the paper and/or the Supplementary Materials. Additional data related to this paper may be requested from the authors.

## Notes

### Competing Interest Statement

The authors have declared no competing interest.

## References

Abu-Hassan, D.W., Acott, T.S., Kelley, M.J., 2014. The Trabecular Meshwork: A Basic Review of Form and Function. J Ocul Biol 2, 9.

Acott, T.S., Kelley, M.J., 2008. Extracellular matrix in the trabecular meshwork. Exp Eye Res 86, 543–561.

Agarwal, P., Daher, A.M., Agarwal, R., 2015. Aqueous humor TGF-β2 levels in patients with open-angle glaucoma: A meta-analysis. Molecular vision 21, 612–620.

Alexander, J.P., Samples, J.R., Acott, T.S., 1998. Growth factor and cytokine modulation of trabecular meshwork matrix metalloproteinase and TIMP expression. Curr Eye Res 17, 276–285.

Deravi, L.F., Sinatra, N.R., Chantre, C.O., Nesmith, A.P., Yuan, H., Deravi, S.K., Goss, J.A., MacQueen, L.A., Badrossamy, M.R., Gonzalez, G.M., Phillips, M.D., Parker, K.K., 2017. Design and Fabrication of Fibrous Nanomaterials Using Pull Spinning. Macromolecular Materials and Engineering 302, 1600404.

Drury, J.L., Mooney, D.J., 2003. Hydrogels for tissue engineering: scaffold design variables and applications. Biomaterials 24, 4337–4351.

Fleenor, D.L., Shepard, A.R., Hellberg, P.E., Jacobson, N., Pang, I.H., Clark, A.F., 2006. TGFbeta2-induced changes in human trabecular meshwork: implications for intraocular pressure. Invest Ophthalmol Vis Sci 47, 226–234.

Fuchshofer, R., Tamm, E.R., 2009. Modulation of extracellular matrix turnover in the trabecular meshwork. Experimental Eye Research 88, 683–688.

Fuchshofer, R., Tamm, E.R., 2012. The role of TGF-β in the pathogenesis of primary open-angle glaucoma. Cell Tissue Res 347, 279–290.

Fuchshofer, R., Welge-Lussen, U., Lütjen-Drecoll, E., 2003. The effect of TGF-beta2 on human trabecular meshwork extracellular proteolytic system. Exp Eye Res 77, 757–765.

Granstein, R.D., Staszewski, R., Knisely, T.L., Zeira, E., Nazareno, R., Latina, M., Albert, D.M., 1990. Aqueous humor contains transforming growth factor-beta and a small (less than 3500 daltons) inhibitor of thymocyte proliferation. J Immunol 144, 3021–3027.

Green, J.J., Elisseeff, J.H., 2016. Mimicking biological functionality with polymers for biomedical applications. Nature 540, 386–394.

Han, H., Wecker, T., Grehn, F., Schlunck, G., 2011. Elasticity-Dependent Modulation of TGF-β Responses in Human Trabecular Meshwork Cells. Investigative Ophthalmology & Visual Science 52, 2889–2896.

Hann, C.R., Fautsch, M.P., 2011. The elastin fiber system between and adjacent to collector channels in the human juxtacanalicular tissue. Invest. Ophthalmol. Vis. Sci. 52, 45–50.

Hann, C.R., Springett, M.J., Wang, X., Johnson, D.H., 2001. Ultrastructural localization of collagen IV, fibronectin, and laminin in the trabecular meshwork of normal and glaucomatous eyes. Ophthalmic Res 33, 314–324.

Hensel, N., Rademacher, S., Claus, P., 2015. Chatting with the neighbors: crosstalk between Rho-kinase (ROCK) and other signaling pathways for treatment of neurological disorders. Frontiers in Neuroscience 9.

Honjo, M., Tanihara, H., Inatani, M., Kido, N., Sawamura, T., Yue, B.Y., Narumiya, S., Honda, Y., 2001. Effects of rho-associated protein kinase inhibitor Y-27632 on intraocular pressure and outflow facility. Invest Ophthalmol Vis Sci 42, 137–144.

Inatani, M., Tanihara, H., Katsuta, H., Honjo, M., Kido, N., Honda, Y., 2001. Transforming growth factor-beta 2 levels in aqueous humor of glaucomatous eyes. Graefes Arch Clin Exp Ophthalmol 239, 109–113.

Jensen, C., Teng, Y., 2020. Is It Time to Start Transitioning From 2D to 3D Cell Culture? Frontiers in Molecular Biosciences 7.

Kasetti, R.B., Maddineni, P., Patel, P.D., Searby, C., Sheffield, V.C., Zode, G.S., 2018. Transforming growth factor β2 (TGFβ2) signaling plays a key role in glucocorticoid-induced ocular hypertension. J Biol Chem 293, 9854–9868.

Keller, K.E., Acott, T.S., 2013. The Juxtacanalicular Region of Ocular Trabecular Meshwork: A Tissue with a Unique Extracellular Matrix and Specialized Function. J Ocul Biol 1, 3.

Keller, K.E., Bhattacharya, S.K., Borras, T., Brunner, T.M., Chansangpetch, S., Clark, A.F., Dismuke, W.M., Du, Y., Elliott, M.H., Ethier, C.R., Faralli, J.A., Freddo, T.F., Fuchshofer, R., Giovingo, M., Gong, H., Gonzalez, P., Huang, A., Johnstone, M.A., Kaufman, P.L., Kelley, M.J., Knepper, P.A., Kopczynski, C.C., Kuchtey, J.G., Kuchtey, R.W., Kuehn, M.H., Lieberman, R.L., Lin, S.C., Liton, P., Liu, Y., Lutjen-Drecoll, E., Mao, W., Masis-Solano, M., McDonnell, F., McDowell, C.M., Overby, D.R., Pattabiraman, P.P., Raghunathan, V.K., Rao, P.V., Rhee, D.J., Chowdhury, U.R., Russell, P., Samples, J.R., Schwartz, D., Stubbs, E.B., Tamm, E.R., Tan, J.C., Toris, C.B., Torrejon, K.Y., Vranka, J.A., Wirtz, M.K., Yorio, T., Zhang, J., Zode, G.S., Fautsch, M.P., Peters, D.M., Acott, T.S., Stamer, W.D., 2018. Consensus recommendations for trabecular meshwork cell isolation, characterization and culture. Exp Eye Res 171, 164–173.

Kelley, M.J., Rose, A.Y., Keller, K.E., Hessle, H., Samples, J.R., Acott, T.S., 2009. Stem cells in the trabecular meshwork: present and future promises. Exp Eye Res 88, 747–751.

Khademhosseini, A., Langer, R., 2007. Microengineered hydrogels for tissue engineering. Biomaterials 28, 5087–5092.

Lampi, M.C., Reinhart-King, C.A., 2018. Targeting extracellular matrix stiffness to attenuate disease: From molecular mechanisms to clinical trials. Sci Transl Med 10.

Lee, K.Y., Mooney, D.J., 2001. Hydrogels for tissue engineering. Chem. Rev. 101, 1869–1879.

Li, H., Bagué, T., Kirschner, A., Strat, A.N., Roberts, H., Weisenthal, R.W., Patteson, A.E., Annabi, N., Stamer, W.D., Ganapathy, P.S., Herberg, S., 2021. A tissue-engineered human trabecular meshwork hydrogel for advanced glaucoma disease modeling. Exp Eye Res 205, 108472.

Liu, J., Wada, Y., Katsura, M., Tozawa, H., Erwin, N., Kapron, C.M., Bao, G., Liu, J., 2018. Rho-Associated Coiled-Coil Kinase (ROCK) in Molecular Regulation of Angiogenesis. Theranostics 8, 6053–6069.

Liu, Y., Suzuki, Y.J., Day, R.M., Fanburg, B.L., 2004. Rho Kinase–Induced Nuclear Translocation of ERK1/ERK2 in Smooth Muscle Cell Mitogenesis Caused by Serotonin. Circulation Research 95, 579–586.

Ma, J., Sanchez-Duffhues, G., Goumans, M.-J., Ten Dijke, P., 2020. TGF-β-Induced Endothelial to Mesenchymal Transition in Disease and Tissue Engineering. Front Cell Dev Biol 8, 260–260.

Medina-Ortiz, W.E., Belmares, R., Neubauer, S., Wordinger, R.J., Clark, A.F., 2013. Cellular fibronectin expression in human trabecular meshwork and induction by transforming growth factor-β2. Invest Ophthalmol Vis Sci 54, 6779–6788.

Ochiai, Y., Ochiai, H., 2002. Higher concentration of transforming growth factor-beta in aqueous humor of glaucomatous eyes and diabetic eyes. Jpn J Ophthalmol 46, 249–253.

Pattabiraman, P.P., Rao, P.V., 2010. Mechanistic basis of Rho GTPase-induced extracellular matrix synthesis in trabecular meshwork cells. Am J Physiol Cell Physiol 298, C749–763.

Picht, G., Welge-Luessen, U., Grehn, F., Lutjen-Drecoll, E., 2001. Transforming growth factor beta 2 levels in the aqueous humor in different types of glaucoma and the relation to filtering bleb development. Graefes Arch Clin Exp Ophthalmol 239, 199–207.

Prendes, M.A., Harris, A., Wirostko, B.M., Gerber, A.L., Siesky, B., 2013. The role of transforming growth factor β in glaucoma and the therapeutic implications. Br J Ophthalmol 97, 680–686.

Quigley, H.A., 1993. Open-angle glaucoma. N Engl J Med 328, 1097–1106.

Raghunathan, V.K., Morgan, J.T., Dreier, B., Reilly, C.M., Thomasy, S.M., Wood, J.A., Ly, I., Tuyen, B.C., Hughbanks, M., Murphy, C.J., Russell, P., 2013. Role of substratum stiffness in modulating genes associated with extracellular matrix and mechanotransducers YAP and TAZ. Invest Ophthalmol Vis Sci 54, 378–386.

Ramachandran, C., Patil, R.V., Combrink, K., Sharif, N.A., Srinivas, S.P., 2011. Rho-Rho kinase pathway in the actomyosin contraction and cell-matrix adhesion in immortalized human trabecular meshwork cells. Molecular vision 17, 1877–1890.

Rao, P.V., Deng, P.F., Kumar, J., Epstein, D.L., 2001. Modulation of aqueous humor outflow facility by the Rho kinase-specific inhibitor Y-27632. Invest Ophthalmol Vis Sci 42, 1029–1037.

Rao, P.V., Pattabiraman, P.P., Kopczynski, C., 2017. Role of the Rho GTPase/Rho kinase signaling pathway in pathogenesis and treatment of glaucoma: Bench to bedside research. Exp. Eye Res. 158, 23–32.

Ren, R., Li, G., Le, T.D., Kopczynski, C., Stamer, W.D., Gong, H., 2016. Netarsudil Increases Outflow Facility in Human Eyes Through Multiple Mechanisms. Invest Ophthalmol Vis Sci 57, 6197–6209.

Seliktar, D., 2012. Designing cell-compatible hydrogels for biomedical applications. Science 336, 1124–1128.

Serini, G., Bochaton-Piallat, M.L., Ropraz, P., Geinoz, A., Borsi, L., Zardi, L., Gabbiani, G., 1998. The fibronectin domain ED-A is crucial for myofibroblastic phenotype induction by transforming growth factor-beta1. The Journal of cell biology 142, 873–881.

Serle, J.B., Katz, L.J., McLaurin, E., Heah, T., Ramirez-Davis, N., Usner, D.W., Novack, G.D., Kopczynski, C.C., 2018. Two Phase 3 Clinical Trials Comparing the Safety and Efficacy of Netarsudil to Timolol in Patients With Elevated Intraocular Pressure: Rho Kinase Elevated IOP Treatment Trial 1 and 2 (ROCKET-1 and ROCKET-2). Am J Ophthalmol 186, 116–127.

Sethi, A., Jain, A., Zode, G.S., Wordinger, R.J., Clark, A.F., 2011. Role of TGFbeta/Smad signaling in gremlin induction of human trabecular meshwork extracellular matrix proteins. Invest Ophthalmol Vis Sci 52, 5251–5259.

Stamer, W.D., Clark, A.F., 2017. The many faces of the trabecular meshwork cell. Exp Eye Res 158, 112–123.

Stamer, W.D., Seftor, R.E., Williams, S.K., Samaha, H.A., Snyder, R.W., 1995. Isolation and culture of human trabecular meshwork cells by extracellular matrix digestion. Curr. Eye Res. 14, 611–617.

Sun, K.-H., Chang, Y., Reed, N.I., Sheppard, D., 2016. α-Smooth muscle actin is an inconsistent marker of fibroblasts responsible for force-dependent TGFβ activation or collagen production across multiple models of organ fibrosis. American Journal of Physiology-Lung Cellular and Molecular Physiology 310, L824–L836.

Tamm, E.R., 2009. The trabecular meshwork outflow pathways: structural and functional aspects. Exp Eye Res 88, 648–655.

Tamm, E.R., Russell, P., Epstein, D.L., Johnson, D.H., Piatigorsky, J., 1999. Modulation of myocilin/TIGR expression in human trabecular meshwork. Invest Ophthalmol Vis Sci 40, 2577–2582.

Tanna, A.P., Johnson, M., 2018. Rho Kinase Inhibitors as a Novel Treatment for Glaucoma and Ocular Hypertension. Ophthalmology 125, 1741–1756.

Tong, J., Li, L., Ballermann, B., Wang, Z., 2016. Phosphorylation and Activation of RhoA by ERK in Response to Epidermal Growth Factor Stimulation. PLoS One 11, e0147103.

Tovar-Vidales, T., Clark, A.F., Wordinger, R.J., 2011. Transforming growth factor-beta2 utilizes the canonical Smad-signaling pathway to regulate tissue transglutaminase expression in human trabecular meshwork cells. Exp Eye Res 93, 442–451.

Wang, G., McCain, M.L., Yang, L., He, A., Pasqualini, F.S., Agarwal, A., Yuan, H., Jiang, D., Zhang, D., Zangi, L., Geva, J., Roberts, A.E., Ma, Q., Ding, J., Chen, J., Wang, D.Z., Li, K., Wang, J., Wanders, R.J., Kulik, W., Vaz, F.M., Laflamme, M.A., Murry, C.E., Chien, K.R., Kelley, R.I., Church, G.M., Parker, K.K., Pu, W.T., 2014. Modeling the mitochondrial cardiomyopathy of Barth syndrome with induced pluripotent stem cell and heart-on-chip technologies. Nat Med 20, 616–623.

Wang, J., Liu, X., Zhong, Y., 2013. Rho/Rho-associated kinase pathway in glaucoma (Review). Int. J. Oncol. 43, 1357–1367.

Wang, S.K., Chang, R.T., 2014. An emerging treatment option for glaucoma: Rho kinase inhibitors. Clin. Ophthalmol. 8, 883–890.

Welge-Lüßen, U., May, C.A., Lütjen-Drecoll, E., 2000. Induction of Tissue Transglutaminase in the Trabecular Meshwork by TGF-β1 and TGF-β2. Investigative Ophthalmology & Visual Science 41, 2229–2238.

Zhang, K., Zhang, L., Weinreb, R.N., 2012. Ophthalmic drug discovery: novel targets and mechanisms for retinal diseases and glaucoma. Nat Rev Drug Discov 11, 541–559.

Zhang, Y.E., 2009. Non-Smad pathways in TGF-β signaling. Cell Research 19, 128–139.

Zhang, Y.S., Khademhosseini, A., 2017. Advances in engineering hydrogels. Science 356, eaaf3627.

Zhao, X., Ramsey, K.E., Stephan, D.A., Russell, P., 2004. Gene and Protein Expression Changes in Human Trabecular Meshwork Cells Treated with Transforming Growth Factor-β. Investigative Ophthalmology & Visual Science 45, 4023–4034.

Zhao, Y., Lv, M., Lin, H., Hong, Y., Yang, F., Sun, Y., Guo, Y., Cui, Y., Li, S., Gao, Y., 2012. ROCK1 induces ERK nuclear translocation in PDGF-BB-stimulated migration of rat vascular smooth muscle cells. IUBMB Life 64, 194–202.

