## Supplemental information for "TGFβ2 regulates human trabecular meshwork cell contractility via ERK and ROCK pathways with distinct signaling crosstalk dependent on the culture substrate"

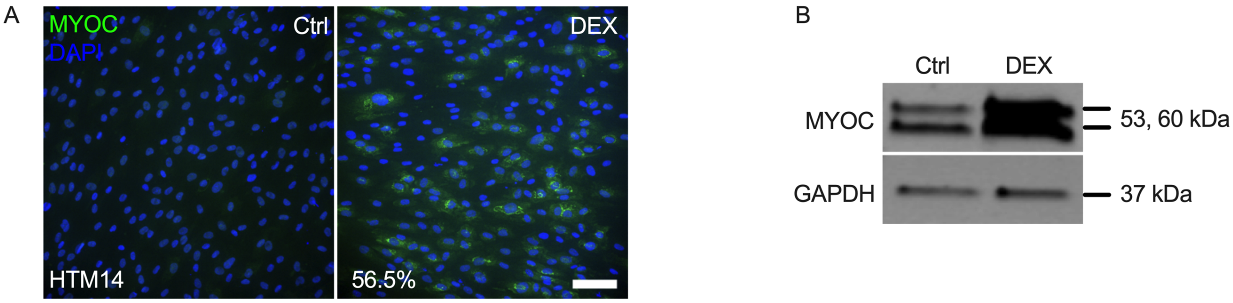


**Suppl. Fig. 1. HTM14 cell characterization.** (A) Representative fluorescence micrographs of intracellular MYOC at 7 d (MYOC = green; DAPI = blue). Scale bar, 100 µm. (B) Immunoblot of secreted MYOC at 7 d.

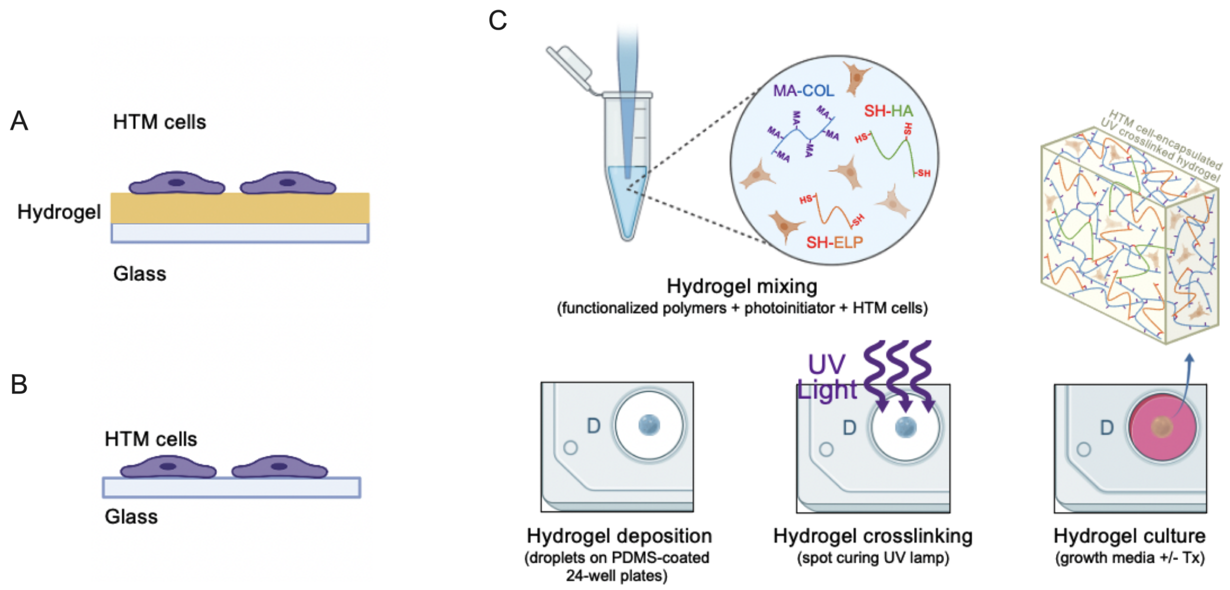


**Suppl. Fig. 2. Schematic of sample preparation.** (A and B) HTM cells were seeded on pre-made hydrogels or conventional glass coverslips. (C) HTM cell-laden hydrogel preparation. Created with BioRender.com.

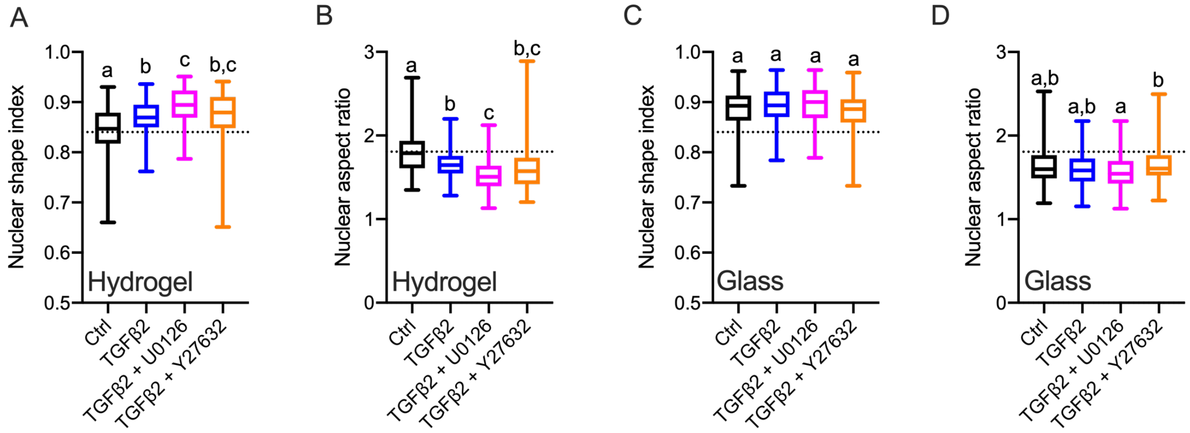


**Suppl. Fig. 3. Effects of TGFβ2 in absence or presence of ERK or ROCK inhibition on nuclear shape index and aspect ratio.** (A,C) Nuclear shape index and (B,D) nuclear aspect ratio of HTM cells on hydrogels and glass subjected to the different treatments at 3 d (n = 100 nuclei from three biological replicates; dotted lines show mean value of control HTM cells on hydrogel for reference). The box and whisker plots represent median values (horizontal bars), 25th to 75th percentiles (box edges) and minimum to maximum values (whiskers); shared significance indicator letters represent non-significant difference ($p>0.05$), distinct letters represent significant difference ($p<0.05$).

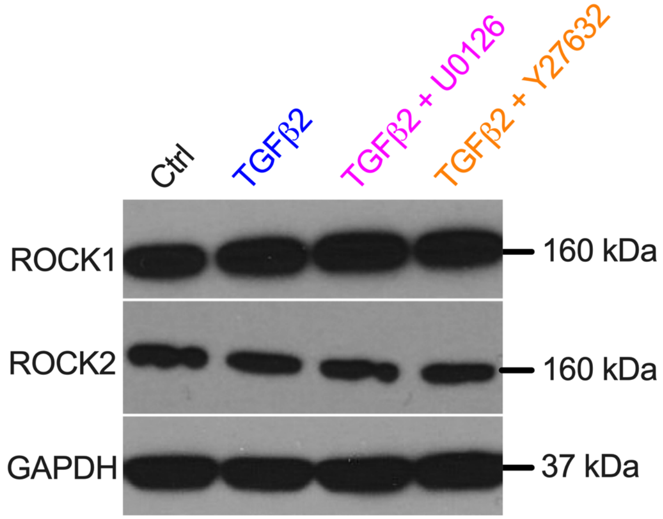


| | | | | |
|---|---|---|---|---|
| ROCK1/ GAPDH | 1 | 1.25 | 1.45 | 1.20 |
| ROCK2/ GAPDH | 1 | 0.92 | 0.90 | 0.90 |

**Suppl. Fig. 4. Effects of TGFβ2 in absence or presence of ERK or ROCK inhibition on ROCK1 and ROCK2 expression.** Immunoblots of ROCK1, ROCK2, and GAPDH with relative fold-changes vs. control.
